# A pseudo-meiotic centrosomal function of TEX12 in cancer

**DOI:** 10.1101/509869

**Authors:** S Sandhu, LJ Salmon, JE Hunter, CL Wilson, ND Perkins, N Hunter, OR Davies, UL McClurg

**Affiliations:** Howard Hughes Medical Institute, Department of Microbiology and Molecular Genetics, University of California, Davis, CA 95616, USA.; Institute for Cell and Molecular Biosciences, Newcastle University, Newcastle upon Tyne NE2 4HH, UK.; Institute of Cellular Medicine, Newcastle University, Newcastle upon Tyne NE2 4HH, UK.; Institute for Integrative Biology, University of Liverpool, Liverpool, L69 7ZB, UK.

## Abstract

Cell division by meiosis involves an extraordinary chromosome choreography including pairing, synapsis and crossing over between homologous chromosomes^1, 2^. The many meiosis-specific genes involved in these processes also constitute a latent toolbox of chromosome remodelling and recombination factors that may be exploited through aberrant expression in cancer^3, 4^. Here, we report that TEX12, a structural protein involved in meiotic chromosome synapsis^5–7^, is aberrantly expressed in human cancers, with high TEX12 levels correlating with poor prognosis. We find that TEX12 knock-down causes proliferative failure in multiple cancer cell lines, and confirm its role in the early stages of oncogenesis through murine cancer models. Remarkably, somatically expressed TEX12 localises to centrosomes, leading to altered centrosome number and structure, features associated with cancer development. Further, we identify TEX12 in meiotic centrin-rich bodies, likely precursors of the mitotic centrosome, suggesting that this may represent an additional cellular function of TEX12 in meiosis that has been previously overlooked. Thus, we propose that an otherwise meiotic function of TEX12 in centrosome duplication is responsible for promoting oncogenesis and cellular proliferation in cancer, which may be targeted for novel cancer therapeutics and diagnostics.

## Main text

The process of reductive cell division by meiosis is characterised by the assembly of the synaptonemal complex (SC)^1, 8^. This supramolecular protein structure binds together homologous chromosome pairs and is essential for the maturation of inter-homologue crossover recombination events, and thereby fertilty^1, 2, 8, 9^. The structural roles of the SC include chromosome remodelling through axis compaction, chromatin loop stabilisation by SYCP3^10^, and the physical tethering of homologues by an SYCP1 zipper-like lattice that is stabilised by a series of SC central element proteins^11^. The central element includes a 4:4 complex between TEX12 and SYCE2, which undergoes filament-like self-assembly into structures that are thought to enable SC growth along the entire length of meiotic chromosomes^5–7^. On the basis of their roles in chromosome structure, and their predicted functional interaction with the DNA recombination machinery, we reasoned that SC components could be detrimental to chromosomal stability if expressed outside the regulated process of meiosis, and thus could contribute to the onset and/or maintenance of cancer. In support of this, SYCP1 and central element component SYCE1 have been reported as cancer testis antigens (CTA)^12^ a defined group of genes which instead of being present only in male germ cells can be abberantly re-expressed in cancer causing an immune reaction. Furthermore, SYCP3 expression has been identified in a range of primary tumours, with a proposed role in suppressing BRCA2-mediated inter-sister recombination^13^.

To assess the role of SC proteins in human cancer, we utilised a large-scale transcriptomic analysis of ovarian cancer patient material^14^ to determine SC protein expression and its correlation with disease severity. We identified the presence of multiple SC proteins in ovarian cancer patients with their expression levels correlating with survival (data not shown). This was most striking in the case of TEX12, where its amplification correlated with a remarkably poor prognosis indicated by the absence of any long-term survivors in the TEX12-amplified cohort (Fig. 1a). We found a similar correlation of TEX12 amplification with poor survival in a number of other cancers, including glioblastoma^15^ (Fig. 1b), kidney renal clear cell carcinoma, thyroid carcinoma and uterine corpus endometrial carcinoma but not in all cases as TEX12 expression levels are unrelated to survival in prostate cancer^16^ (Fig. 1c). These findings suggest a specific pro-oncogenic functions of TEX12 in a subgroup of cancers.

**Figure 1.**
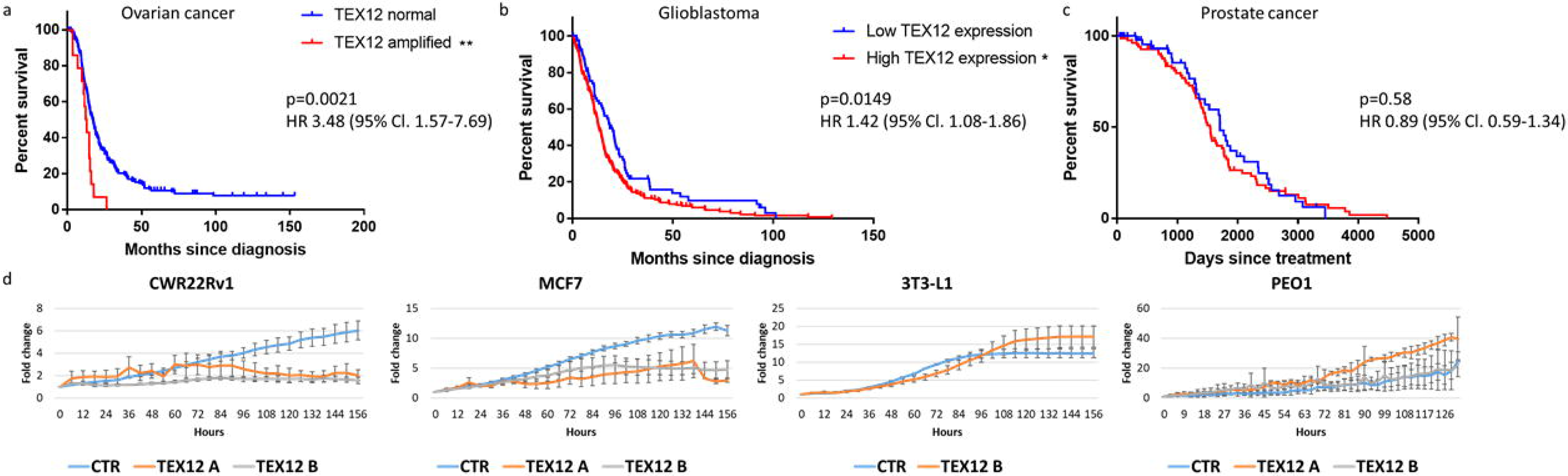
TEX12 is a marker of poor prognosis in cancers and is required for cancer cell proliferation. (**a**) Kaplan-Meier curve of TCGA dataset of ovarian cancer. Patients were divided based on presence and absence of TEX12 amplification. (**b**) Kaplan-Meier survival curve of TCGA dataset of glioblastoma. Patients were divided based on high versus low TEX12 transcripts. (**c**) Kaplan-Meier survival curve of TCGA dataset of prostate cancer. Patients were divided based on high versus low TEX12 transcripts. (**d**) Cells were treated with siRNA and cell growth was measured every 3h with IncuCyte. *p<0.05; **p<0.001; ***p<0.005.

We first assessed how TEX12 expression might lead to poor prognosis in human cancers. To test whether TEX12 is important for cancer cell growth, we performed siRNA knockdown of TEX12 in multiple cancer cell lines (Fig. 1d and Supplementary Fig. 1a-b). In the majority of cancer cell lines tested, TEX12 silencing abrogated cellular proliferation, with a failure to increase cell numbers when cultured for a period of up to seven days (Fig. 1d). In contrast, TEX12-negative PEO1 cancer cells and non-transformed fibroblasts showed no response to TEX12 silencing (Fig. 1d and Supplementary Fig. 1b). We further confirmed that this proliferative defect was not simply due to cell death (Supplementary Fig. 1c), establishing that TEX12 is required for cell proliferation in a subset of human cancers.

We next examined the role of TEX12 in tumourigenesis using mouse models of lymphoma and hepatocellular carcinoma (Fig. 2a-c). In an Eµ-myc model of MYC-driven B-cell lymphoma^17^, Tex12 protein was absent in the mesenteric lymph nodes of naïve mice while it was highly expressed in the lymph nodes of all lymphoma-positive mice, as assessed by immunohistochemistry (Fig. 2a). Further, high lymph node expression of Tex12 was correlated with a more aggressive disease and shortened survival (Fig. 2b), analogous to our observations of TEX12 expression in human ovarian and glioblastoma cancers. We also detected Tex12 expression in a diethylnitrosamine (DEN)-induced mouse model of hepatocellular carcinoma^18, 19^ (Fig. 2c), indicating that these correlations are not limited to lymphoma or MYC-driven tumours. Importantly, Tex12 expression was detected as early as five weeks following DEN treatment, with peak levels observed at 40 weeks (Fig. 2c). Its detection at the five week-stage indicates that Tex12 is already expressed at the very early stages of oncogenesis, prior to the formation of an overt tumour. This observation raises the possibility that TEX12 is a driver of early oncogenesis and highlights its potential as an early diagnostic marker.

**Figure 2.**
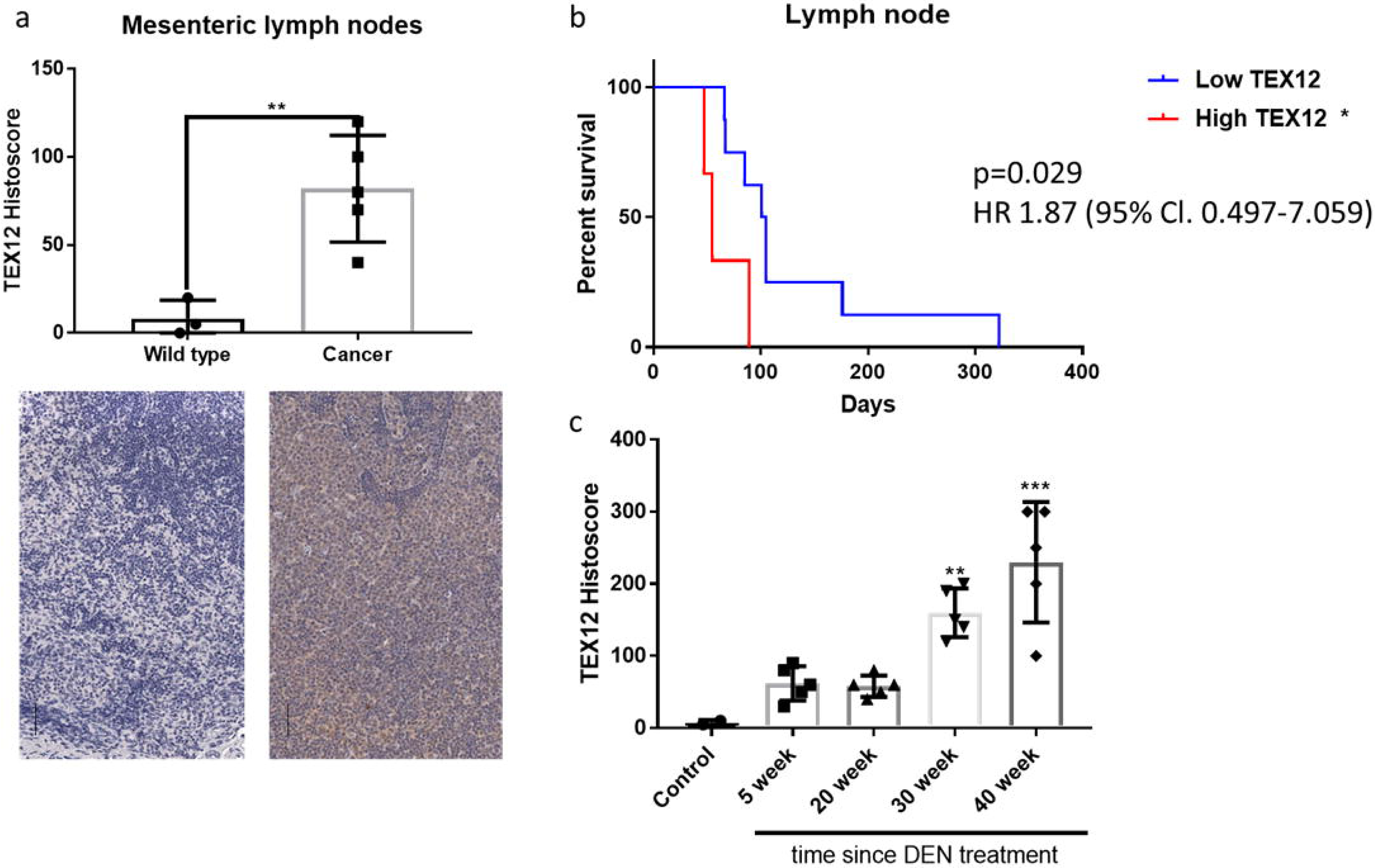
TEX12 is expressed with the onset of cancer. (**a**) Eµ-Myc lymphoma and healthy controls mesenteric lymph nodes were stained and scored for Tex12 proteinlevels. Each data point indicates one animal. Black lines indicate 50µm. **p<0.01. (**b**) Kaplan-Meier survival curve of Eµ-myc lymphoma mice divided based on their lymph node Tex12 protein histoscore. (**c**) Mice have been scored for liver Tex12 protein levels following DEN-treatment tumour induction. Each data point represents one animal.

To determine how TEX12 could contribute to cancer cell proliferation we performed immunofluorescent staining in a number of TEX12-expressing cancer cell lines using commercial TEX12 antibody (Supplementary Fig. 2 A-C), consistently revealing the presence of TEX12 in dot-like peri-nuclear foci (Fig. 3a). The number and location of TEX12 foci resembled that of centrosomes. Accordingly, we found that TEX12 co-localised with both centrin and pericentrin foci (Fig. 3b,c), confirming its recruitment to centrosomes. Furthermore, FLAG-tagged TEX12 expressed in TEX12-negative COS7 cells also co-localised with pericentrin in dot-like foci (Fig. 3d). Thus, we conclude that TEX12 is recruited to centrosomes in cancer cells.

**Figure 3.**
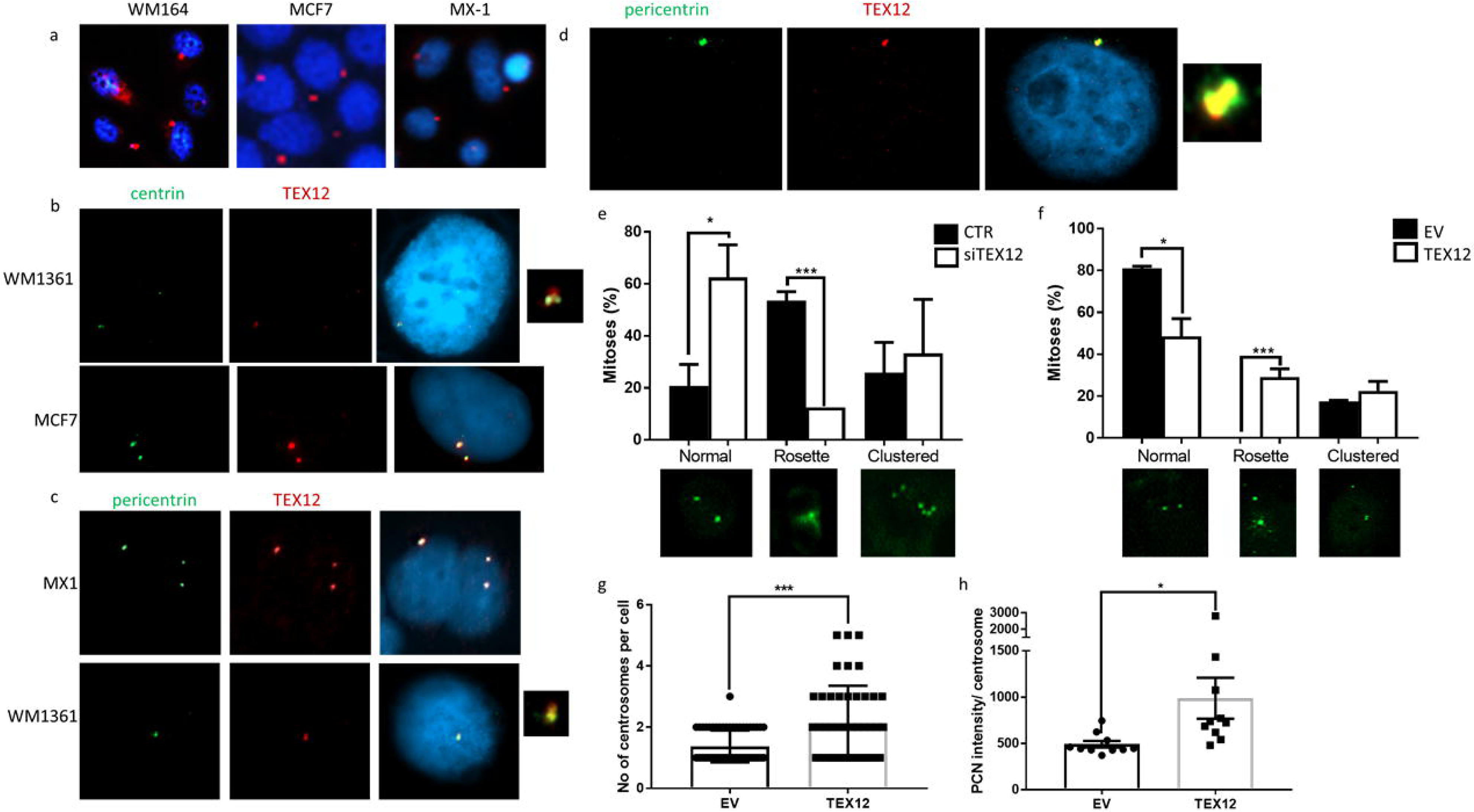
TEX12 is a centrosomal protein and regulates centrosome division in cancer. (**a**) Cells were stained for TEX12 visible in red with DAPI visible in blue. (**b**) Cells were stained for TEX12 (red), centrin (green) and nuclei were visualised in blue with DAPI. (**c**) Cells were stained for TEX12 (red), pericentrin (green) and nuclei were visualised in blue with DAPI. (**d**) COS-7 cells were transfected with TEX12 for 48h following by staining for TEX12 (red) and pericentrin (green). (**e**) Cancer cells were treated with non silencing control (CTR) siRNA or siTEX12 for 96h followed by centrin staining in green. Mitosis types were quantified. *p<0.05; ***p<0.005. (**f**) COS-7 cells were transfected with either empty vector or TEX12 for 96h followed by centrin staining in green. Mitosis types were quantified. *p<0.05; ***p<0.005. (**g**) Centrosome numbers in COS-7 cells 96h after transfection with TEX12 or empty vector were counted. ***p<0.005. (**h**) COS-7 cells were transfected with empty vector control (EV) or TEX12 for 96h followed by TEX12 and pericentrin imaging. Average pericentrin intensity per centrosome was quantified. *p<0.05.

Interestingly, we observed that TEX12 expression is also associated with a diffuse centrosomal staining pattern indicative of rosette centrosomes, which have recently been described as a common feature of cancer cells (Fig. 3e, f)^20–22^. Rosette centrosomes consist of mature centrosomes surrounded by a rosette of immature daughter centrioles, formed upon aberrant overduplication of centrioles, and are associated with biased chromosome capture and chromosome missegregation owing to asymmetric force of uneven number of microtubules bound to daughter centrioles at two spindle poles^22–24^. The number of cancer cells with rosette centrosomes was substantially reduced upon TEX12 silencing (Fig. 3e), and similarly the number of rosette centrosomes positive cells was increased upon expression of TEX12 in TEX12-negative COS7 cells (Fig. 3f). Further, the overall number of centrosomes and pericentrin intensity were increased upon TEX12 expression (Fig. 3g-h). Thus, in addition to its recruitment to centrosomes in cancer cells, TEX12 actively contributes towards their amplification and structural stability.

In meiosis, TEX12 assembles into a 4:4 oligomeric complex with SYCE2 to form a structural component of the SC^5–7^. Thus, we addressed whether SYCE2, which we also found expressed in some cancer cells, is similarly recruited to centrosomes. Immunofluorescent staining of SYCE2 in cancer cells revealed a predominantly cytoplasmic pattern with no dot-like foci or overt co-localisation with pericentrin, indicating that centrosome-associated TEX12 is not in complex with SYCE2 (Fig. 4a). However, co-expression of SYCE2 and TEX12 in COS7 cells, a non-cancer cell line normally negative for both TEX12 and SYCE2, did lead to SYCE2 recruitment to centrosomes (Fig. 4b,c). Thus, whilst TEX12 can be recruited to centrosomes in absence of SYCE2 (Fig. 3f-h), centrosome-associated TEX12 retains its SYCE2-binding ability and SYCE2 is recruited to centrosomes only when co-expressed with TEX12.

**Figure 4.**
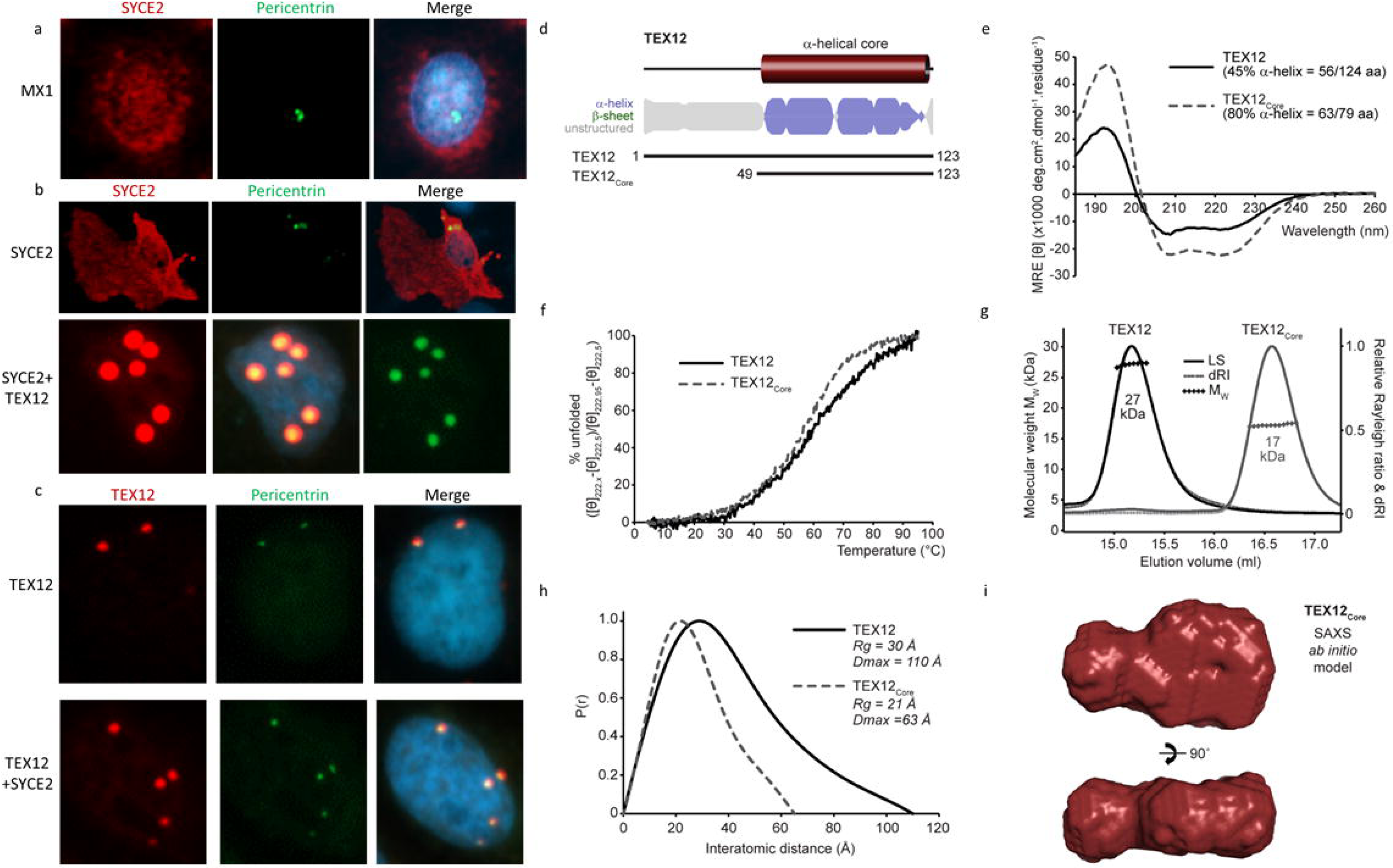
TEX12 forms a stable α-helical dimer and can localize to centrosomes independently of its SC partner, SYCE2. (**a**) MX1 breast cancer cells were stained for SYCE2 (red) and pericentrin (green). Nuclei were visualised in blue with DAPI. (**b**) COS-7 cells were transfected with SYCE2 and TEX12 as indicated. Cells were stained for SYCE2 (red) and pericentrin (green). Nuclei were visualised in blue with DAPI. (**c**) COS-7 cells were transfected with SYCE2 and TEX12 as indicated. Cells were stained for TEX12 (red) and pericentrin (green). Nuclei were visualised in blue with DAPI. (**d**) Sequence analysis of human TEX12 demonstrating the presence of a core α-helical region in the C-terminal half of the protein. Secondary structure prediction is shown as per residue scores and protein constructs are indicated along with their sequence boundaries. (**e**) Far UV CD spectra and (**f**) CD thermal denaturation of TEX12 (solid line) and TEX12core (dashed line). (**e**) Secondary structure composition was estimated through deconvolution of spectra with data fitted at normalised r.m.s. deviation values of 0.015 and 0.008, respectively. (**f**) Thermal denaturation was recorded as % unfolded based on the helical signal at 222 nm; melting temperatures were estimated at 60°C and 57°C, respectively. (**g**) SEC-MALS analysis of TEX12 and TEX12core in which light scattering (LS) and differential refractive index (dRI) are overlaid, with and fitted molecular weights (Mw) plotted as diamonds across elution peaks. TEX12 and TEX12core are both dimeric, with molecular weights of 27 kDa and 17 kDa, respectively (theoretical dimer sizes - 28 kDa and 18 kDa). (**h-i**) SEC-SAXS analysis of TEX12 and TEX12core. (**h**) P(r) distributions of TEX12 (solid line) and TEX12core (dashed line) showing maximum dimensions of 110 Å and 63 Å, respectively. (**i**) SAXS ab initio model of the TEX12core dimer. A filtered averaged model was generated from 30 independent DAMMIF runs, with NSD=0.609 (± 0.047) and reference model χ2=1.11.

We aimed to explain the ability of TEX12 to function in centrosomes independent of its binding partner SYCE2. Whilst recombinant co-expression of human SYCE2 and TEX12 was essential for the biochemical stability of SYCE2 *in vitro*, TEX12 was soluble and stable in absence of SYCE2 (Fig. 4d and Supplementary Fig. 3a, b). Analysis of the secondary structural composition, stability and oligomeric state of TEX12 by CD and SEC-MALS revealed that it is a largely α-helical homodimer, formed by a structural core of C-terminal amino-acids 49-123 (Fig. 4e-g), which corresponds to its SYCE2-bindinging region^7^. Further, analysis of protein size and shape by SEC-SAXS determined that the structural core adopts a largely globular conformation, with dimensions consistent with it forming a flattened four-helical structure of two helix-loop-helix chains (Fig. 4h-i and Supplementary Fig. 3c), similar to the reported structure of another SC protein, SYCE3^25^. The specific dimeric structure adopted by TEX12 provides important validation for its SYCE2-independent centrosome function.

The pathological effect of TEX12 on centrosome function in cancer cells raises the possibility that its physiological function during meiosis might also include a centrosome function, in addition to its well-established role in chromosome synapsis. While it is generally thought that centrosomes dissipate in gametogenesis, with the loss of centrioles and formation of asymmetric microtubule organising centres, recent studies have reported the retention of centriole remnants in mature oocytes^26–30^. In mice it has been extensively demonstrated that oocytes contribute the centrosomes while centrosomal proteins and centrioles was reported to come from the sperm^31^. Moreover, in a published analysis of Tex12 localisation in SC-defective *Sycp1* mutant spermatocytes^5^ we noticed the presence of one or two prominent Tex12 immunostaining foci, and wondered if these might be centrosome-related structures. Therefore, we immune-stained wild-type mouse spermatocytes for Tex12 and the core centriole marker Centrin (Fig 5a). Close juxtaposition of Tex12 and Centrin was observed throughout meiotic prophase I and metaphase I (MI; Fig 5a i-viii). In prophase I, TEX12 was localized peripheral to nuclei as an elongated or bi-lobed structure, two closely apposed foci, or two more widely separated foci. Centrin was localized immediately adjacent to these Tex12 structures, typically appearing as three or four distinct foci, indicating that centrosomes have been duplicated. During diakinesis/metaphase-I, Tex12 structures morphed into two internal toroids, each associated with two Centrin foci atop these structures (Fig 5a vi-viii). In some metaphase-I nuclei, the two Tex12-Centrin structures appeared quite elongated and positioned on either side of the chromosome mass, with the Centrin foci on the outer, poleward faces (Fig 5a viii).

**Figure 5.**
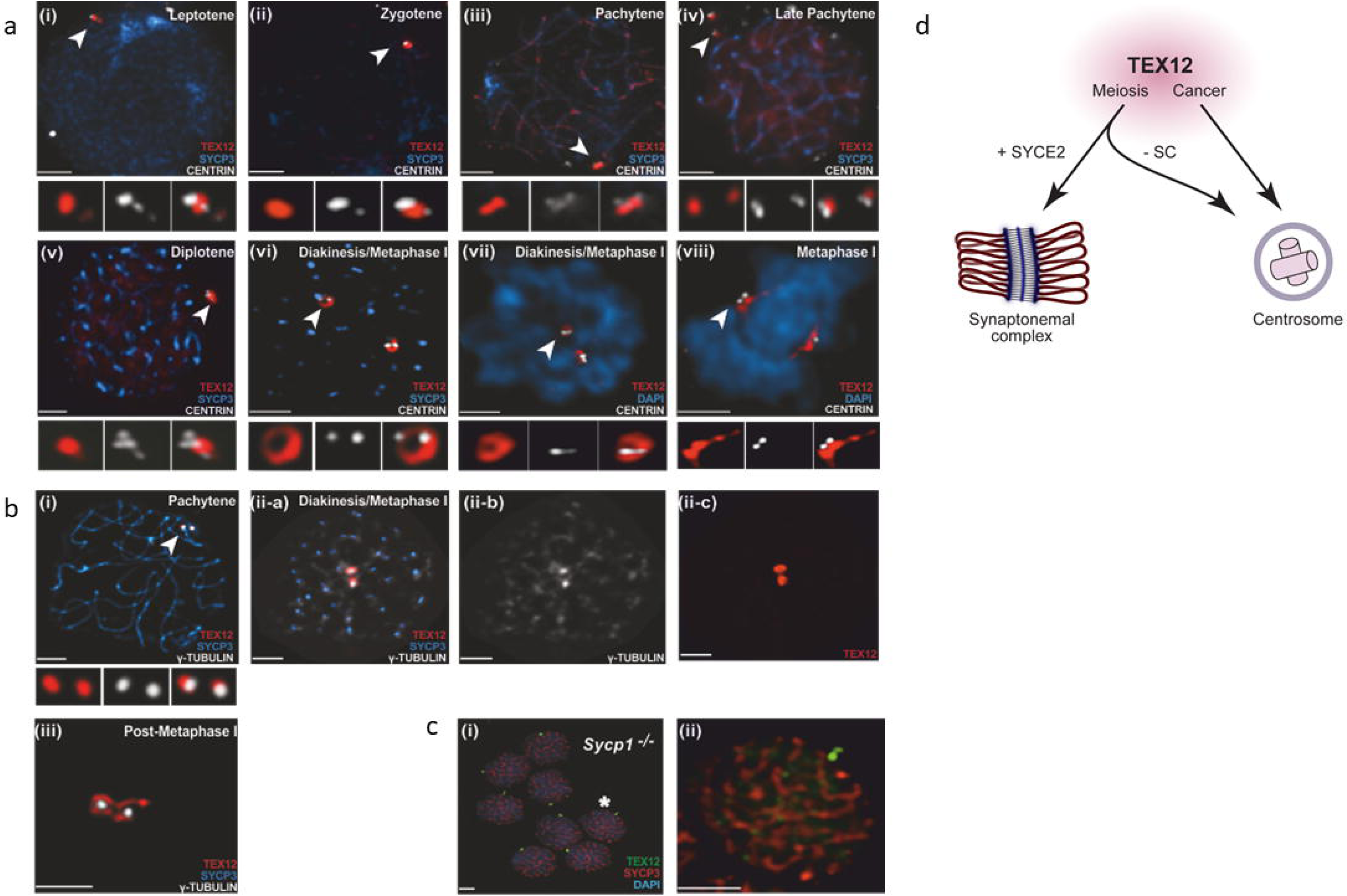
TEX12 is expressed in centrin rich bodies in meiosis. (**a**) (i-vi) Mouse spermatocyte squashes stained for Centrin in blue, Sycp3 in green and Tex12 in red. (**a**) (vii-viii) Mouse metaphase I spermatocyte squashes stained for DAPI in blue, Centrin in green and Tex12 in red. (**b**) Mouse spermatocyte squashes stained for Sycp3 in blue, Tex12 in red and γ-tubulin in white. (**c**) Mouse spermatocyte squashes from Sycp1 knockout animals stained for DAPI in blue, Sycp3 in red and Tex12 in green. (**d**) Model of TEX12 activity.

Tex12 was also localized relative to the microtubule organizing centre (MTOC) marker, γ-tubulin (Fig 5b). In prophase-I spermatocytes, the two Tex12 foci partially overlapped with bright γ-tubulin signals (Fig 5b i). During diakinesis/metaphase-I, the two Tex12 toroids appeared to encircle two brightly staining γ-tubulin structures (Fig 5b ii). Numerous additional γ-tubulin structures appeared at this time, many of which localized to centromere regions (that retained residual Sycp3 protein) and presumably nucleate kinetochore-associated microtubules (Fig 5b ii). In post MI spermatocytes (that had lost residual Sycp3), two bright γ-tubulin-staining MTOCs remained, which were closely apposed and surrounded by interconnected Tex12 toroids (Fig 5b iii). These structures are presumptively the precursors of MII spindles.

Finally, we confirmed that Tex12 still localized to centrosomal structures in the *Sycp1* mutant (Fig 5c), suggesting that its role at centrosomes is independent of its function in the SC. Taken together, our localization studies are consistent with TEX12 being a meiosis-specific component of the outer pericentriolar material (PCM) where it may augment or replace another PCM factor to facilitate meiotic spindle function. Our data suggest that TEX12 that does not get incorporated into SCs may be recruited to these Centrin-rich bodies, where it may perform a similar function to that observed in centrosomes of cancer cells, supporting centrosomal division. This likely represents a role of TEX12 that is favoured when it cannot participate in SC assembly, which may be analogous to the formation of chromatin-free SYCP1 polycomplexes in the absence of an SC in meiotic and somatic cells^32, 33^. We therefore propose that, in keeping with the mechanism of other meiosis-specific genes in cancer^3, 4^, the centrosomal role of TEX12 in cancer cells may represent a previously overlooked pseudo-meiotic function.

Here, we report that TEX12, a structural component of the meiotic SC, is expressed in cancer at centrosomes and is indicative of poor prognosis. This dichotomy of TEX12 localisation resembles previous observations regarding Sme4/PaMe4 in fungal sexual cycle where it first takes part in the SC formation and then, after disappearing and reappearing, localises to the spindle pole body^34^. The localisation and function of TEX12 in centrosomes provides a crucial specificity that contradicts the previous rationale that the cancer transcriptome is more promiscuous and meiotic gene reexpression is a simple byproduct of cancer cell biology^35^. Further, the identification of a similar localisation pattern in mouse meiosis indicates that the role of TEX12 in centrosomes reflects a re-activation of a function that may be important for the regulation of TEX12 and SC levels, and/or for the role of centrosome-like bodies in meiosis (Fig 5d). Additionally, it has previously been shown that centrosome amplification is sufficient to initiate tumourigenesis^36, 37^, with centrosome amplification observed in the majority of cancers^38^. Thus, the role of TEX12 in centrosome division and rosette formation provides an attractive explanation for its involvement in the early stages of oncogenesis and correlation with poor prognosis in human cancers. Furthermore, it has been recently reported that supernumerary centrosomes induce the IL-8 paracrine-signalling axis in breast cancer, leading to increased migration and invasion of cells with amplified centrosomes into the wider cellular population. This could be especially important in tumours with TEX12 amplification, such as aggressive ovarian tumours (Fig 1a), as chromosomal *TEX12* is proximal to *IL-8* and hence both genes could easily be amplified together^39^. We propose that TEX12 constitutes a specific prognostic marker and a target for novel therapeutics and diagnostics in a number of human cancers.

## Materials and methods

### Antibodies and plasmids

The anti-TEX12, anti-pericentrin and anti-SYCE2 antibodies were obtained from Abcam and anti-FLAG antibody was purchased from Sigma. SYCE2 and TEX12 were cloned and expressed using FLAG-pCMV-HA vector.

### Cell culture, transfections, and lentiviral transductions

LNCaP and PC3 prostate cancer, MCF7 and MX1 breast cancer, WM1361 NRAS mutant and WM164 BRAF mutant melanoma, 3T3-L1 mouse fibroblasts and COS-7 cells were obtained from American Type Culture Collection (Manassas, USA). LNCaP, PC3 and MX-1 cells were maintained in RPMI 1640 media and remaining cell lines in DMEM media supplemented with 2mM L-glutamine (Invitrogen) and 10% (v/v) fetal calf serum (FCS) at 37°C in 5% CO_2_. Cell lines were never maintained for more than 30 passages or 2 months of continuous culturing. Cell lines were tested for mycoplasma on a tri-monthly basis. Proliferation was measured by live cell imaging with the Incucyte system every 6 hours for 156 hours post-treatment. Transfections were performed using Lipofectamine2000 reagent (Invitrogen) following the manufacturer’s instructions.

### siRNA gene silencing and gene expression analysis

The *TEX12* targeting siRNA sequences were (A) AAGCCUUGGAGAAAGAUUUAAAU[dTdT], (B) CAGCAGUAGAUGCAUCUUACA[dTdT] and (C) SASI_Hs01_00169803 (Sigma). Cells were reverse transfected with siRNA using RNAiMax (Invitrogen) according to manufacturer’s instructions and incubated in culture media for 96 hours prior to cell lysis and analysis. For real-time qPCR, total RNA was extracted using TRIzol (Invitrogen, 15596-026), RNA quality and yields were assessed using a NanoDrop 2000 (NanoDrop), 1μg of total RNA was reverse transcribed using SuperScript VILO (Invitrogen, 11755-050), and qPCR performed using QuantiTect SYBR Green (QIAGEN, 204143) on an ABI PRISM 7500 Sequence Detection System (Applied Biosystems). Data was tested for parametric distribution. Parametric data was analysed using appropriate t-tests or ANOVA with Bonferroni’s comparison test for multiple group comparisons. Non-parametric data was analysed using Wilcoxon signed-rank test. By convention, p-values < 0.001 are marked with ***, < 0.01 with **, and < 0.05 with *.

### Immunohistochemistry

Antigens were retrieved by microwaving the slides in 10mM citrate pH 6.0 for 15 minutes followed by staining the tissues. Antibodies were detected with ImmPRESS HRP IgG (Vector labs). Samples were scored blind using the histoscore methodology^40^. Briefly, percentage and intensity of staining for positive cells was estimated (0, 1, 2, 3) using the following equation: H-score = (% of cells with low level positivity) + 2 × (% of cells with medium level positivity) + 3 × (% of cells with high level positivity). For survival analysis, high marker levels were defined as a value in the third and fourth upper quarter of the population

### Immunofluorescence

For Immunofluorescence, cells were fixed for 10 min in ice cold methanol followed by blocking in 1% BSA, staining and imaging using confocal microscope.

### Recombinant protein expression and purification

Human TEX12 sequences, corresponding to amino acids 1-123 (full length) and 49-123 (core), were cloned into the pMAT11 vector (Peranen et al., 1996) for bacterial expression as a fusion protein to an N-terminal His_6_-MBP-tag, separated by a tobacco etch virus (TEV) site for tag cleavage. Proteins were expressed in BL21(DE3) E. coli cells (Novagen®), cultured in 2xYT media and induced with 0.5 mM IPTG at 25° for 16 hours. Cells were lysed in 20 mM Tris pH 8.0, 500 mM KCl, by sonication and clarified lysate was applied to Ni-NTA resin (Qiagen) followed by amylose resin (NEB). Proteins were further purified by TEV protease cleavage for tag removal, anion exchange (GE Healthcare) and stored at -80°C in 20 mM Tris pH 8.0, 150 mM KCl, 2 mM DTT after concentration (PALL Microsep™ Advance, 3kD) to 6-12 mg/ml. Proteins were analysed by SDS-PAGE (NuPAGE Bis-Tris system) with SimplyBlue SafeStain (Invitrogen). UV spectrophotometry (Cary 60 UV-Vis, Agilent Technologies) was used to calculate protein concentrations with theoretical molecular weights and extinction coefficients determined by ExPASY ProtParam.

### Circular dichroism (CD) spectroscopy

CD data were collected using a Jasco J-810 spectropolarimeter (Institute for Cell and Molecular Biosciences, Newcastle University). CD spectra for TEX12 and TEX12_Core_ were collected in 10 mM NaH_2_PO_4_ pH 7.5, 150 mM NaF, at 0.36 and 0.24 mg/ml respectively. Measurements were recorded at 4°C between 260 and 185 nm, with 0.2 nm intervals, in a 0.2 mm pathlength quartz cuvette (Hellma). Nine measurements were collected and averaged, with buffer correction, and converted to mean residue ellipticity (MRE [θ]). The DichroWeb server (http://dichroweb.cryst.bbk.ac.uk) was used for used for deconvolution with the CDSSTR algorithm. CD thermal melts were recorded in 20 mM Tris pH 8.0, 150 mM KCl, 2 mM DTT, at 222 nm with a temperature range of 5-95°C and a 1°C per minute ramping rate. Data were plotted as % unfolded after conversion to MRE ([θ]_222,x_-[θ]_222,5_)/([θ]_222,95_-[θ]_222,5_) with melting temperatures (Tm) determined by the temperature at which the sample is 50 % unfolded.

### Size-exclusion chromatography multi-angle light scattering (SEC-MALS)

SEC-MALS was performed to determine the absolute molecular masses of TEX12 and TEX12_Core_. Protein samples, at 12 and 4 mg/ml respectively, were loaded onto a pre-equilibrated (20 mM Tris pH 8.0, 150 mM KCl, 2mM DTT) Superdex™ 200 Increase 10/300 GL SEC column (GE Healthcare) with an ÄKTA™ Pure (GE Healthcare) at 0.5 ml/min. The column flow-through was directed to a DAWN HELEOS II MALS detector (Wyatt Technology) and then to an Optilab T-rEX (Wyatt Technology) differential refractometer. ASTRA^®^ 6 software (Wyatt Technology) was used calculate molecular weights from the eluted peaks, by Zimm plot extrapolation using a 0.1850 ml/g dn/dc value.

### Size-exclusion chromatography small-angle X-ray scattering (SEC-SAXS)

SEC-SAXS data collection was carried out at the B21 beamline at the Diamond Light Source synchrotron facility (Oxfordshire, UK). TEX12 and TEX12_Core_, at 12 and 6 mg/ml respectively, were loaded onto a pre-equilibrated (20 mM Tris pH 8.0, 150 mM KCl, 2mM DTT) Superdex™ 200 Increase 10/300 GL SEC column (GE Healthcare) with an Agilent 1200 HPLC system at 0.5 ml/min. The column flow-through was directed to the experimental cell for SAXS data collection in 3 s frames, with a 4.014 m detector distance at 12.4 keV. ScÅtter 3.0 (http://www.bioisis.net) was used to analyse SAXS data. Data were subtracted for buffer, and then averaged for Guinier (Rg) and cross-sectional radius (Rc) analysis. *P(r)* distributions were generated using PRIMUS^41^ and *ab initio* modelling was performed using DAMMIF^42^.

Ethics statement - All animal experiments were approved by Newcastle University’s Animal Welfare and Ethical Review Board and all procedures were carried out under project and personal licences approved by the Secretary of State for the Home Office, under the Animals in Scientific Procedures Act (1986).

### Mouse models

Eμ-Myc mice^17^ were purchased from The Jackson Laboratory, Maine, USA and wild-type (C57Bl/6) were purchased from Charles River, UK. Colonies were established and maintained in the Comparative Biology Centre, Newcastle University, according to the FELASA Guidelines. No blinding of groups in mouse studies was performed. All mice were designated to an experimental group dependent on their genotype. To perform survival analysis, Eμ-Myc transgenic mice were monitored daily and were sacrificed at pre-determined end-points, defined as the animal becoming moribund, losing bodyweight/condition and/or having palpable tumour burden at any lymphoid organ site. At this point, mice were necropsied and samples taken for downstream analysis.

### Spermatocyte staining

Testis were dissected from freshly sacrificed mice and processed for cell squashes as previously described^43^. Briefly, seminiferous tubules were fixed in freshly prepared 2% formaldehyde containing 0.1% Triton-X for 10 mins at room temperature. Small pieces of tubules were placed on a glass slide, minced gently and then squashed under a coverslip with pressure from the blunt end of pencil. Slides were then frozen briefly in liquid nitrogen and coverslips removed. Following three washes with PBS, immunofluorescent staining was performed as described^44^, using the following primary antibodies with incubation overnight at room temperature: rabbit anti-TEX12 (Abcam, ab122455;1:200 dilution), mouse anti-Centrin (Millipore, 04-1624; 1:200 dilution), mouse anti-SYCP3-488 (Abcam, ab205846; 1:100 dilution), mouse anti-γ-Tubulin-647 (Abcam; 1:100 dilution). Goat secondary antibodies were then added for 1 hour at 37°C, either anti-rabbit 488 (Life Technologies, 31822; 1:1000 dilution), or anti-mouse 594 (Life Technologies, A-11032, 1:1000 dilution). Finally, slides were rinsed, stained with DAPI and mounted with ProLong Gold antifade reagent. Images were acquired using a Zeiss AxioPlan II microscope, 63X, 1.4 NA objective and X-Cite light source. Images were captured by a Hamamatsu ORCA-ER CCD camera and processed using Volocity (PerkinElmer).

## Acknowledgements

ULM was supported by the Newcastle’s Fellowship Programme, in part funded through the Wellcome Trust’s Institutional Strategic Support Fund followed by the IIB Fellowship programme and by the Royal Society Research Grant RG170342. We thank Diamond Light Source and the staff of beamline B21 (proposals sm15897, sm15580 and sm15836), and H. Waller for assistance with CD data collection. ORD is a Sir Henry Dale Fellow jointly funded by the Wellcome Trust and Royal Society (Grant Number 104158/Z/14/Z) and is also supported by Royal Society Research Grant RG170118. SS was funded by the A.P. Giannini Postdoctoral Research Fellowship.

## Author Contributions

Conceptualisation: ULM and ORD; Investigation: ULM, SS, LIS and JH; Mass spectrometry: MWP; Biochemical analysis: LJS; Meiotic investigation: SS and NH; Animal work: CLW, JEH, NDP, IHC: ULM; Writing - Original Draft: ULM and ORD; Writing - Review & Editing: ULM and ORD; Funding Acquisition: ULM, NH and ORD

## Supplementary Figure Legends

<u>Supplementary Figure 1 TEX12 is critical for cancer cell proliferation.</u> (**a**) MX1 cancer cells were treated with siRNA as indicated for 96 hours and after RNA extraction TEX12 transcript levels were measured with quantitative RT-PCR and normalised to HPRT1 transcript levels. (**b**) Cells were treated with siRNA and cell growth was measured every 3h with IncuCyte. (**c**) Cells were treated with siRNA for 156h and cell death was measured with CellTox.

<u>Supplementary Figure 2 Antibody validation.</u> (**a**) COS7 cells were transfected with empty vector control plasmid (EV) or TEX12 for 48h followed by immunofluorescence with TEX12 antibodies. (**b**) MX-1 cells were treated with control siRNA (CTR) or siTEX12 for 96h followed by immunofluorescence with TEX12 antibody. (**c**) COS7 cells were transfected with empty vector control or TEX12 for 96h followed by cell lysis and Western blotting with TEX12 antibody.

<u>Supplementary Figure 3 Purification and SAXS analysis of recombinant TEX12.</u> (**a-b**) SDS-PAGE of recombinant expression and purification of (**a**) TEX12 (amino acids 1-123) and (**b**) TEX12Core (amino acids 49-123) through Ni-NTA, amylose, and anion exchange chromatography, following by TEV cleavage to remove N-terminal MBP tags, with subsequent anion exchange and concentration. (**c-d**) SEC-SAXS analysis of TEX12 and TEX12core. (**c**) Scattering intensity plots; experimental data are shown in open circles with P(r) distribution fits displayed as red lines. (**d**) Guinier analysis to determine the radius of gyration (Rg) of TEX12 and TEX12core with linear fits shown in black. Q.Rg values were < 1.3.

## References

1. Hunter, N. Meiotic Recombination: The Essence of Heredity. Cold Spring Harbor perspectives in biology 7 (2015).

2. Baudat, F., Imai, Y. & de Massy, B. Meiotic recombination in mammals: localization and regulation. Nat Rev Genet 14, 794–806 (2013).

3. McFarlane, R.J. & Wakeman, J.A. Meiosis-like Functions in Oncogenesis: A New View of Cancer. Cancer Res (2017).

4. Nielsen, A.Y. & Gjerstorff, M.F. Ectopic Expression of Testis Germ Cell Proteins in Cancer and Its Potential Role in Genomic Instability. Int J Mol Sci 17 (2016).

5. Hamer, G. et al. Characterization of a novel meiosis-specific protein within the central element of the synaptonemal complex. Journal of Cell Science 119, 4025–4032 (2006).

6. Hamer, G. et al. Progression of meiotic recombination requires structural maturation of the central element of the synaptonemal complex. Journal of cell science 121, 2445–2451 (2008).

7. Davies, O.R., Maman, J.D. & Pellegrini, L. Structural analysis of the human SYCE2–TEX12 complex provides molecular insights into synaptonemal complex assembly. Open Biology 2, 120099 (2012).

8. Zickler, D. & Kleckner, N. Recombination, Pairing, and Synapsis of Homologs during Meiosis. Cold Spring Harbor perspectives in biology 7 (2015).

9. Kouznetsova, A., Benavente, R., Pastink, A. & Hoog, C. Meiosis in mice without a synaptonemal complex. PLoS One 6, e28255 (2011).

10. Syrjanen, J.L., Pellegrini, L. & Davies, O.R. A molecular model for the role of SYCP3 in meiotic chromosome organisation. eLife 3 (2014).

11. Dunce, J.M. et al. Structural basis of meiotic chromosome synapsis through SYCP1 self-assembly. Nat Struct Mol Biol 25, 557–569 (2018).

12. Almeida, L.G. et al. CTdatabase: a knowledge-base of high-throughput and curated data on cancer-testis antigens. Nucleic acids research 37, D816–819 (2009).

13. Hosoya, N. et al. Synaptonemal complex protein SYCP3 impairs mitotic recombination by interfering with BRCA2. EMBO Rep 13, 44–51 (2012).

14. Network, C.G.A.R. Integrated genomic analyses of ovarian carcinoma. Nature 474, 609–615 (2011).

15. Network, C.G.A.R. Comprehensive genomic characterization defines human glioblastoma genes and core pathways. Nature 455, 1061–1068 (2008).

16. Taylor, B.S. et al. Integrative genomic profiling of human prostate cancer. Cancer cell 18, 11–22 (2010).

17. Harris, A.W. et al. The E mu-myc transgenic mouse. A model for high-incidence spontaneous lymphoma and leukemia of early B cells. The Journal of experimental medicine 167, 353–371 (1988).

18. Rajewsky, M.F., Dauber, W. & Frankenberg, H. Liver carcinogenesis by diethylnitrosamine in the rat. Science (New York, N.Y.) 152, 83–85 (1966).

19. Bartsch, H., Malaveille, C. & Montesano, R. In vitro metabolism and microsome-mediated mutagenicity of dialkylnitrosamines in rat, hamster, and mouse tissues. Cancer research 35, 644–651 (1975).

20. Ganem, N.J., Godinho, S.A. & Pellman, D. A mechanism linking extra centrosomes to chromosomal instability. Nature 460, 278–282 (2009).

21. Schnerch, D. & Nigg, E.A. Structural centrosome aberrations favor proliferation by abrogating microtubule-dependent tissue integrity of breast epithelial mammospheres. Oncogene 35, 2711–2722 (2016).

22. Cosenza, M.R. et al. Asymmetric Centriole Numbers at Spindle Poles Cause Chromosome Missegregation in Cancer. Cell reports 20, 1906–1920 (2017).

23. Guderian, G., Westendorf, J., Uldschmid, A. & Nigg, E.A. Plk4 trans-autophosphorylation regulates centriole number by controlling betaTrCP-mediated degradation. Journal of cell science 123, 2163–2169 (2010).

24. Arquint, C., Sonnen, K.F., Stierhof, Y.D. & Nigg, E.A. Cell-cycle-regulated expression of STIL controls centriole number in human cells. Journal of cell science 125, 1342–1352 (2012).

25. Lu, J. et al. Structural insight into the central element assembly of the synaptonemal complex. Scientific reports 4, 7059 (2014).

26. Manandhar, G., Schatten, H. & Sutovsky, P. Centrosome reduction during gametogenesis and its significance. Biology of reproduction 72, 2–13 (2005).

27. Gruss, O.J. Animal Female Meiosis: The Challenges of Eliminating Centrosomes. Cells 7 (2018).

28. Woolley, D.M. & Fawcett, D.W. The degeneration and disappearance of the centrioles during the development of the rat spermatozoon. The Anatomical record 177, 289–301 (1973).

29. Manandhar, G., Simerly, C., Salisbury, J.L. & Schatten, G. Centriole and centrin degeneration during mouse spermiogenesis. Cell motility and the cytoskeleton 43, 137–144 (1999).

30. Simerly, C. et al. Separation and Loss of Centrioles From Primordidal Germ Cells To Mature Oocytes In The Mouse. Scientific reports 8, 12791 (2018).

31. Manandhar, G., Sutovsky, P., Joshi, H.C., Stearns, T. & Schatten, G. Centrosome reduction during mouse spermiogenesis. Developmental biology 203, 424–434 (1998).

32. Goldstein, P. Multiple synaptonemal complexes (polycomplexes): origin, structure and function. Cell Biol Int Rep 11, 759–796 (1987).

33. Ollinger, R., Alsheimer, M. & Benavente, R. Mammalian protein SCP1 forms synaptonemal complex-like structures in the absence of meiotic chromosomes. Mol Biol Cell 16, 212–217 (2005).

34. Espagne, E. et al. Sme4 coiled-coil protein mediates synaptonemal complex assembly, recombinosome relocalization, and spindle pole body morphogenesis. Proceedings of the National Academy of Sciences of the United States of America 108, 10614–10619 (2011).

35. Feichtinger, J., Larcombe, L. & McFarlane, R.J. Meta-analysis of expression of l(3)mbt tumor-associated germline genes supports the model that a soma-to-germline transition is a hallmark of human cancers. International journal of cancer 134, 2359–2365 (2014).

36. Raff, J.W. & Basto, R. Centrosome Amplification and Cancer: A Question of Sufficiency. Developmental cell 40, 217–218 (2017).

37. Godinho, S.A. et al. Oncogene-like induction of cellular invasion from centrosome amplification. Nature 510, 167–171 (2014).

38. Patel, N. et al. Integrated genomics and functional validation identifies malignant cell specific dependencies in triple negative breast cancer. Nature communications 9, 1044 (2018).

39. Arnandis, T. et al. Oxidative Stress in Cells with Extra Centrosomes Drives Non-Cell-Autonomous Invasion. Developmental cell 47, 409–424.e409 (2018).

40. Kirkegaard, T. et al. Observer variation in immunohistochemical analysis of protein expression, time for a change? Histopathology 48, 787–794 (2006).

41. P.V. Konarev, V.V.V., A.V. Sokolova, M.H.J. Koch, D. I. Svergun PRIMUS - a Windows-PC based system for small-angle scattering data analysis. J Appl Cryst. 36, 1277–1282 (2003).

42. Franke, D. & Svergun, D.I. DAMMIF, a program for rapid ab-initio shape determination in small-angle scattering. Journal of applied crystallography 42, 342–346 (2009).

43. Page J, S.J., Santos JL and Rufas JS Squash procedure for protein immunolocalization in meiotic cells. Chromosome Research 6, 639–642 (1998).

44. Reynolds, A. et al. RNF212 is a dosage-sensitive regulator of crossing-over during mammalian meiosis. Nat Genet 45, 269–278 (2013).

