## Supplementary figures and images for "A pseudo-meiotic centrosomal function of TEX12 in cancer"

### Supplemental Figure 1

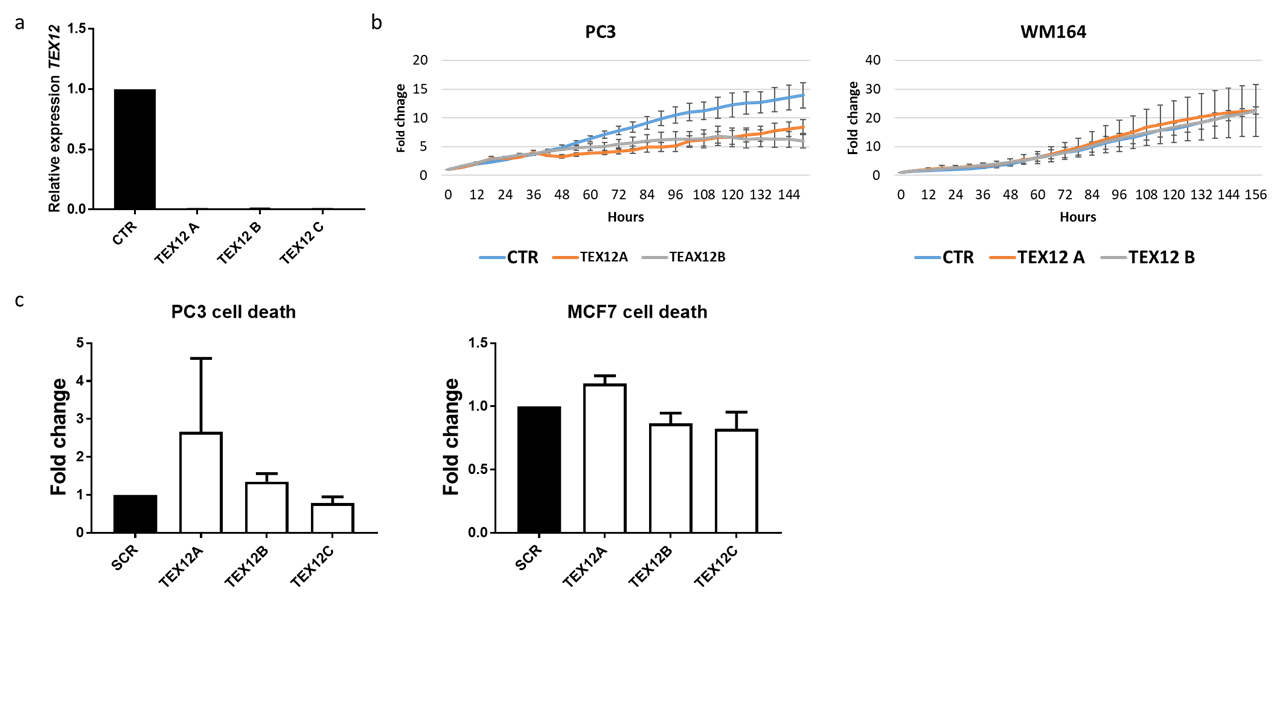

### Supplemental Figure 2

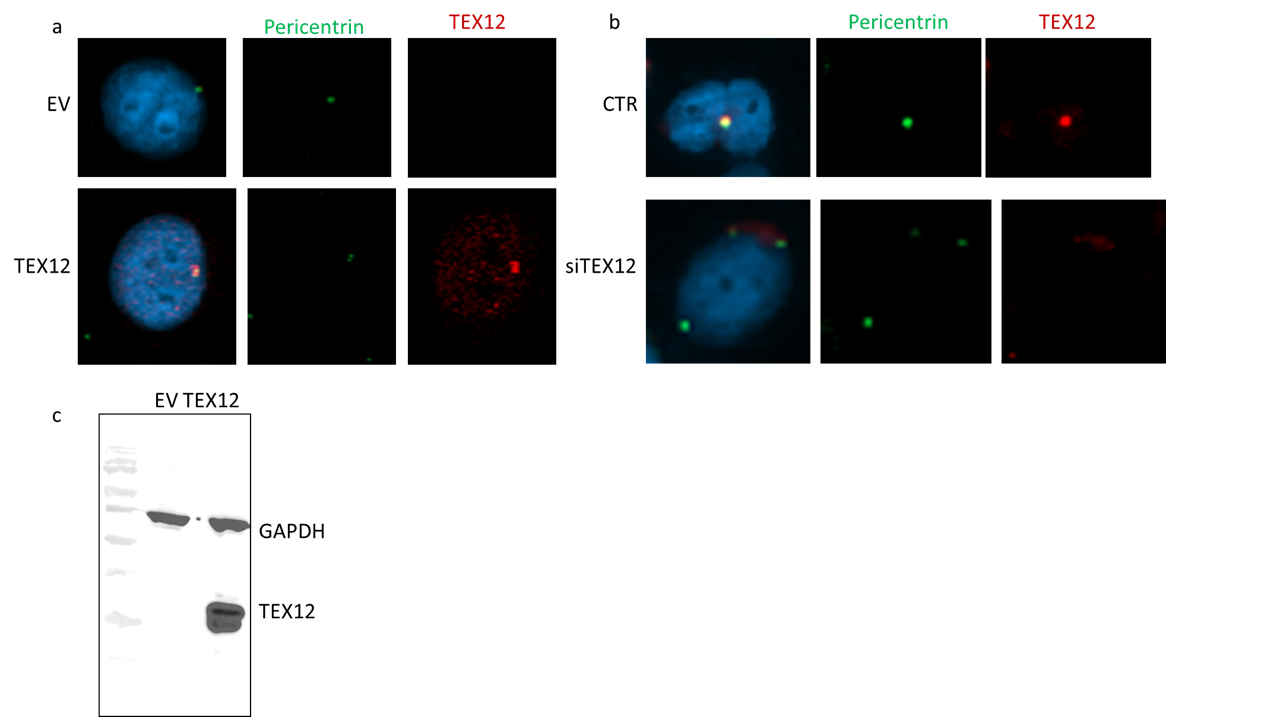

### Supplemental Figure 3

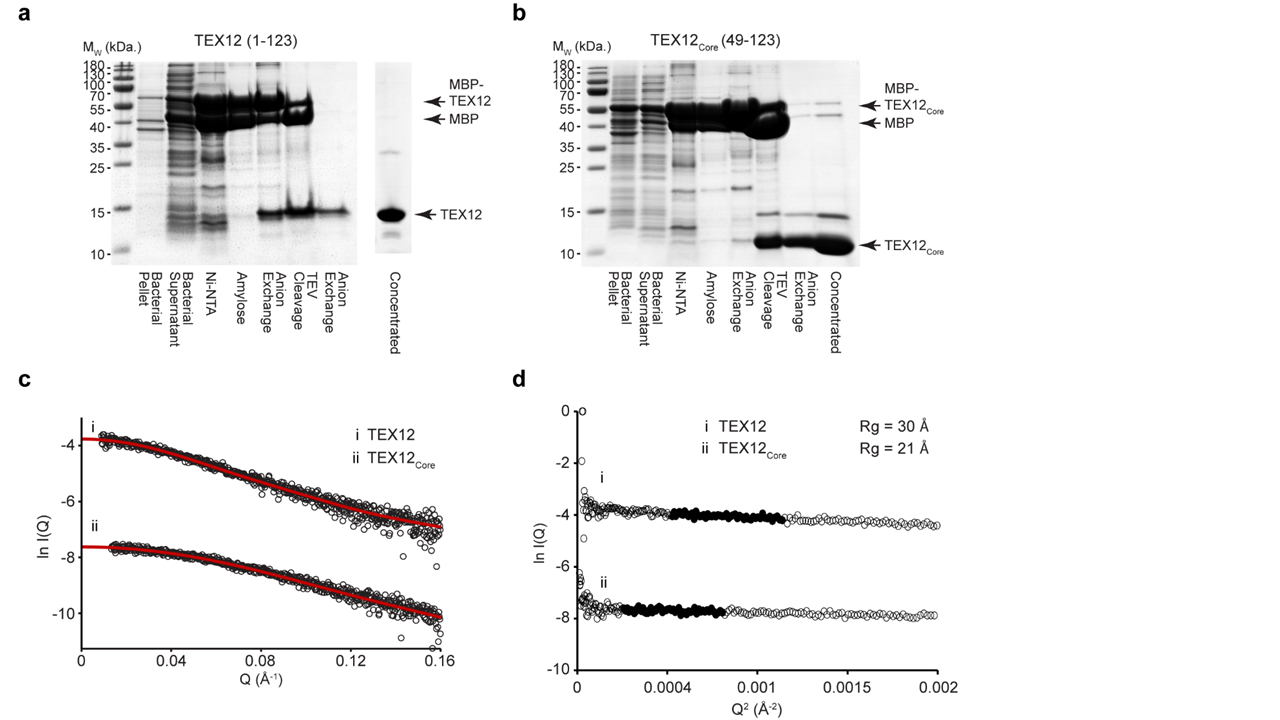
